# A meta-analysis of liver microbiome in beef and dairy cattle using metatranscriptome sequencing reads

**DOI:** 10.1101/2023.03.28.534613

**Authors:** Tansol Park, Wenli Li, Geoffrey Zanton

## Abstract

The gut-liver axis is at the forefront of host-microbial interactions given the proximity of liver to the gut and connection via portal circulation. In recent years, many studies in human and mouse models have demonstrated the existence of a significant microbial community embedded in diverse tissue types, including blood and liver. Yet, in cattle, the rumen microbiome has been the primary focus. The liver microbiome and its metabolic role in host health and performance remain largely unexplored. While there has been considerable work focusing on the liver of diseased cattle, the objective of this study was to evaluate, through meta-analysis, the commensal liver microbiome in various cattle breeds. To our knowledge, this is the first study in which a core liver microbiome has been described in cattle without overt liver disease. We discovered abundant microbial taxa in the liver, varying by host age, species, and developmental stage. Eight bacterial phyla (Actinobacteria, Bacteroidetes, Cyanobacteria, Deinococcus-Thermus, Firmicutes, Fusobacteria, Proteobacteria, and Tenericutes) were found to be the core microbial taxa, representing almost half of the total liver bacterial population. Additionally, we identified several KEGG pathways with significant association with cattle age. This study provides a baseline knowledge of the liver microbiome as identified by metatranscriptome sequencing in cattle. Besides finding the microbial taxa previously reported by studies using DNA-based, 16S rRNA amplicon sequencing methods, this study identified several core phyla that have not been reported in cattle liver, highlighting the improved sensitivity or ability in detecting microbes by RNA-over DNA-based methods.

## Introduction

The gut microbiome is composed of bacteria, protists, fungi, archaea, and viruses naturally residing in the gastrointestinal tract (GIT) with critical roles in several biological processes such as digestion (Henderson et al., 2015), metabolism (Xiao and Kang 2020), and the regulation of the host immune system (Haase, Haghikia et al. 2018). Ruminant animals, including domesticated cattle, sheep, and goats, are especially reliant on maintaining a concentrated microbial community resident in the rumen for converting poorly available energy from their fibrous diet to readily available short-chain fatty acids and microbial protein supporting host productive functions (Wein et al., 2019). For this reason, in beef and dairy cattle, the rumen microbiome has been the primary research focus. However, specific bacteria present in tissues outside of the gut that may be involved in host disease states have also been examined due to the effects on animal health, production, and economics. For instance, *Fusobacterium necrophorum* is considered as the primary etiologic agent of liver abscesses in feedlot cattle through systematic culture studies of the ruminal fluid, rumen lesions, and liver abscesses (Newsom 1938, Madin 1949, Scanlan and Hathcock 1983, Nagaraja and Chengappa 1998, Nagaraja and Lechtenberg 2007). In cattle, the incidence of liver abscesses have been associated with the effects of diet on the ruminal environment and the integrity of the gut epithelium. Diets with a low percentage of roughage (Brink, Lowry et al. 1990) (Harvey, Wise et al. 1968, Gill, Owens et al. 1979, Brink, Lowry et al. 1990), more fermentable grain types, and physical characteristics such as a small particle size (Utley, Hellwig et al. 1973) appear to play important roles in the development of conditions leading to liver abscesses. While the causes and effects of liver abscesses, an overt disease state, have been extensively studied, a general characterization of the liver microbiome and its metabolic role in host health and performance remains largely unexplored.

Recently, many studies in human and mouse subjects have demonstrated the existence of a detectable, and sometimes cultivable, microbial community embedded in diverse tissue types and interior organs, from healthy and diseased tissue. These tissues include adipose (Massier, Chakaroun et al. 2020), brain (Branton, Ellestad et al. 2013), blood (Sato, Kanazawa et al. 2014, Sze, Tsuruta et al. 2014, Mandal, Jiang et al. 2016, Paisse, Valle et al. 2016), and liver (Lluch, Servant et al. 2015, Sookoian, Salatino et al. 2020). Importantly, blood and liver microbial signatures are strongly correlated with various metabolic and digestive diseases (Sato, Kanazawa et al. 2014, Sookoian, Salatino et al. 2020). The liver sits at the junction between the GIT and systemic circulation potentially resulting in translocated GIT bacteria, bacterial products, and their metabolites passing into portal circulation and liver exposure (Henao-Mejia, Elinav et al. 2013, Weinroth, Carlson et al. 2017, Schwenger, Clermont-Dejean et al. 2019). As such, the liver is the final barrier to prevent GIT bacteria and their products, such as LPS, from entering systemic blood circulation (Nanji, Khettry et al. 1994). A unique liver microbial community has been identified in patients with metabolic disorders, including diabetes and non-alcoholic fatty liver syndrome using 16S rRNA gene amplicon sequencing (Iebba, Guerrieri et al. 2018, Anhe, Jensen et al. 2020). Additionally, Anhe and coauthors observed a clear compartmentalization of bacterial communities between tissues such as liver, adipose and blood plasma (Anhe, Jensen et al. 2020).

While these studies provide general support for unique, functional role of liver tissue microbiome in the etiology of liver diseases and disorders in nonruminants, there is little information on the liver microbiome in cattle. Identifying the microbiota in the liver is a first step towards understanding role of this microbiome in cattle and will be transformative for delineating the molecular mechanisms that impact host nutritional and physiological functions. Therefore, the aim of our study was to conduct a meta-analysis on liver whole transcriptome sequence data to identify and characterize the distribution of liver microbiota and associate microbial and functional biomarkers with host phenotype.

## Materials and Methods

### Literature search and study selection

Publicly available, liver whole transcriptome sequencing data were searched from the Sequence Read Archive (SRA) at the National Center for Biotechnology Information (NCBI). Only studies containing all the following information were selected for downstream meta-analysis: species (i.e., *Bos indicus* and *Bos taurus*), breed (i.e., Angus, Brahman, Hereford, Holstein, Jersey, Nellore and Zebu), sex (i.e., male and female), age (i.e., 2 to 63 months). Raw RNA-seqs from those studies were downloaded from NCBI Sequence Read Archive using SRA Toolkit (https://github.com/ncbi/sra-tools) under the following BioProject numbers: PRJEB8831, PRJEB27337, PRJNA263327 (Moran, Cummins et al. 2016), PRJNA303131, PRJNA312148, PRJNA357463 (Salleh, Mazzoni et al. 2017, Salleh, Mazzoni et al. 2018), PRJNA392196, PRJNA393239, PRJNA436715 (Nguyen, Reverter et al. 2018), PRJNA508221, PRJNA526337, PRJNA597813, PRJNA624810 (accessed between 06/2020-07/2020). We also added our unpublished liver RNA-seq data from 2-months-old dairy calves (totally, 8 samples). In total, 251 samples from 14 studies were included in this meta-analysis and sample metadata obtaining from those studies were summarized in Table S1.

**Table 1.** Significant associations with meta-analyzed covariates among minor KEGG-pathways using MaAsLin2.

| Feature | Value | Coefficient | Standard error | No. of samples | No. of samples not zero | P-value | Q-value | Description |  |
| --- | --- | --- | --- | --- | --- | --- | --- | --- | --- |
| Cattle breeds associated |  |  |  |  |  |  |  |  |  |
| ko00196 | Hereford | -0.0151 | 0.0032 | 251 | 44 | 5.14E-06 | 0.0010 | Photosynthesis - antenna proteins | 1.2 Energy metabolism |
| ko00940 | Hereford | -0.0291 | 0.0066 | 251 | 30 | 1.69E-05 | 0.0027 | Phenylpropanoid biosynthesis | 1.10 Biosynthesis of other secondary metabolites |
| ko00941 | Hereford | -0.0269 | 0.0056 | 251 | 43 | 2.53E-06 | 0.0008 | Flavonoid biosynthesis | 1.10 Biosynthesis of other secondary metabolites |
| ko00196 | Holstein | -0.0149 | 0.0038 | 251 | 44 | 0.0001 | 0.0121 | Photosynthesis - antenna proteins | 1.2 Energy metabolism |
| ko00941 | Holstein | -0.0252 | 0.0065 | 251 | 43 | 0.0002 | 0.0153 | Flavonoid biosynthesis | 1.10 Biosynthesis of other secondary metabolites |
| ko04650 | Jersey | 0.0851 | 0.0186 | 251 | 52 | 0.0005 | 0.0410 | Natural killer cell mediated cytotoxicity | 5.1 Immune system |
| Cattle age associated |  |  |  |  |  |  |  |  |  |
| ko00512 | Age | 0.0089 | 0.0016 | 251 | 188 | 1.51E-07 | 6.73E-05 | Mucin type O-glycan biosynthesis | 1.7 Glycan biosynthesis and metabolism |
| ko04062 | Age | -0.0113 | 0.0023 | 251 | 188 | 2.13E-06 | 0.0008 | Chemokine signaling pathway | 5.1 Immune system |
| ko04622 | Age | 0.0051 | 0.0012 | 251 | 140 | 5.27E-05 | 0.0068 | RIG-I-like receptor signaling pathway | 5.1 Immune system |

### Metatranscriptome analysis

The metatranscriptome analysis pipeline was described in our previous study (Park, Cersosimo et al. 2022) with minor modification. Briefly, following the quality checking using FastQC, BBDuk tool (implemented in BBMap v.35.14) was used to filter out low quality reads. High-quality RNA-seq were mapped to the host reference genome UMD 3.1 of *Bos taurus* (http://ccb.jhu.edu/software/tophat/igenomes.shtml, April 1^st^, 2019) using the STAR aligner (Dobin, Davis et al. 2013) to filter out host reads. Active liver-associated microbiota were analyzed using host genome-filtered reads which can be considered as microbial sequences. Bacterial taxonomy classification using Kraken2 (v.2.0.8-beta) (Wood and Salzberg 2014) with a confidence score of 0.2 was done against Kraken2 standard database (built on 4th June 2021). Per million factor normalized read counts per sample was applied to the downstream analysis as described in previous study (Li, Gelsinger et al. 2019, Li, Gelsinger et al. 2019). Bacterial taxa detected in at least 50% of all 251 samples were defined as ‘core liver microbiota’ and mainly discussed in this study.

Microbial functional analysis based on non-rRNA reads separated from Bovine genome-filtered reads by removing rRNA reads (using SortMeRNA (version, 2.1b)) (Kopylova, Noe et al. 2012) with the latest Silva databases (release 138 (Quast, Pruesse et al. 2013)) was done according to the pipeline suggested in Dai et al., 2014 (Dai, Tian et al. 2014). Briefly, non-rRNA reads were blasted against pre-built NCBI nonredundant protein sequence databases (built on May 20th 2020) using blastx module in DIAMOND (Buchfink, Xie et al. 2015) with elevated cutoff of 1e^-5^ and 50 for E-value and bit score, respectively. The deduced amino acid sequences extracted from blastx best-hit were used to search matched KEGG orthologs (KO) using KEGG Automatic Annotation Server (KAAS) (Moriya, Itoh et al. 2007). KEGG pathways were annotated from the KO profiles using the python script implemented in picrust2 package (Douglas, Maffei et al. 2019). KEGG pathways detected at least 90% of all samples were defined as ‘core liver microbial functions’ and mainly discussed in this study.

### Antimicrobial-resistance gene annotation

Active cattle liver microbial resistome was analyzed by aligning the non-rRNA reads against antimicrobial-resistance gene (ARG) databases, MEGARes (version 2.0.0, downloaded on Nov. 20^th^ 2020) (Doster, Lakin et al. 2020) and ResFinder (version 4.0, downloaded on Nov. 20^th^ 2020) (Bortolaia, Kaas et al. 2020), respectively. Non-rRNA reads were blasted (E-value = 1e^-6^, percent identity = 70, query coverage = 60) to the downloaded ARG sequences of those two databases. Blast outputs were normalized and represented in RPKM (Reads per kilo base per million mapped reads) for gene length and sampling depth. Heatmap and hierarchical clustering dendrograms of averaged RPKM of detected ARGs for each ruminant breed were constructed using R package Heatplus (Ploner 2022) based on the Bray-Curtis dissimilarity and using the R function hclust.

### Diversity analysis

Number of identified lowest features (genera for bacteria and KEGG orthologs for microbial functions), evenness, Shannon’s diversity, and Simpson’s diversity indices were analyzed from both the bacterial and KEGG orthologs profiles using vegan (version 2.5-7) R package (Dixon 2003). Compositional abundance of core liver bacteria and microbial functions selected based on the aforementioned occurrence cutoff were used to analyze overall community differences. Principal components analysis was computed to compare the bacterial and microbial functional distribution based on the Bray-Curtis dissimilarity matrices. The R package ggfortify (Tang, Horikoshi et al. 2016) was used to visualize the PCA (principal-component analysis) plots in the two-dimensional space.

### Statistical analysis

The R package MMUPHin was used to reduce the batch effect from different studies while analyzing meta-analysis data (Ma, Shungin et al. 2020). To identify active features associated with all the covariates concurrently available in the metadata (i.e., species, breed, sex, and age), multivariable analysis tool, Microbiome Multivariable Association with Linear Models 2 (MaAsLin2) R package was used (Mallick, Rahnavard et al. 2021). Compositional abundance of core-liver microbiota, core-liver microbial functions, and ARGs were arcsine-transformed but no data transformation was applied for alpha-diversity indices for MaAsLin2 analysis. Only the associations less than 0.05 *Q*-value (Benjamini-Hochberg multiple hypothesis corrected *P*-values) were considered as significant. Permutational multivariate analysis of variance (PERMANOVA) test in vegan (adonis with 9,999 random permutations) was done to statistically identify the overall batch effect adjustment by MMUPHin and the effect of covariates on core-liver microbiome.

## Results

### Core microbiome actively expressed in cattle liver

Following the quality filtering and host genome removal from NCBI SRA downloaded data, an average of 618,673 (standard error of the mean was 74,670) host genome-filtered reads were successfully classified by Kraken2 taxonomic analysis with 21.47% classification rate on average. Eight bacterial phyla (Actinobacteria, Bacteroidetes, Cyanobacteria, Deinococcus-Thermus, Firmicutes, Fusobacteria, Proteobacteria, and Tenericutes) were found to be present in at least half of the analyzed liver samples and representing almost half of the total liver bacterial population (**Figure 1**). Specifically, Proteobacteria and Firmicutes were the first and second dominant bacterial phyla and occupied 47.46% of overall liver microbiota.

**Figure 1.**
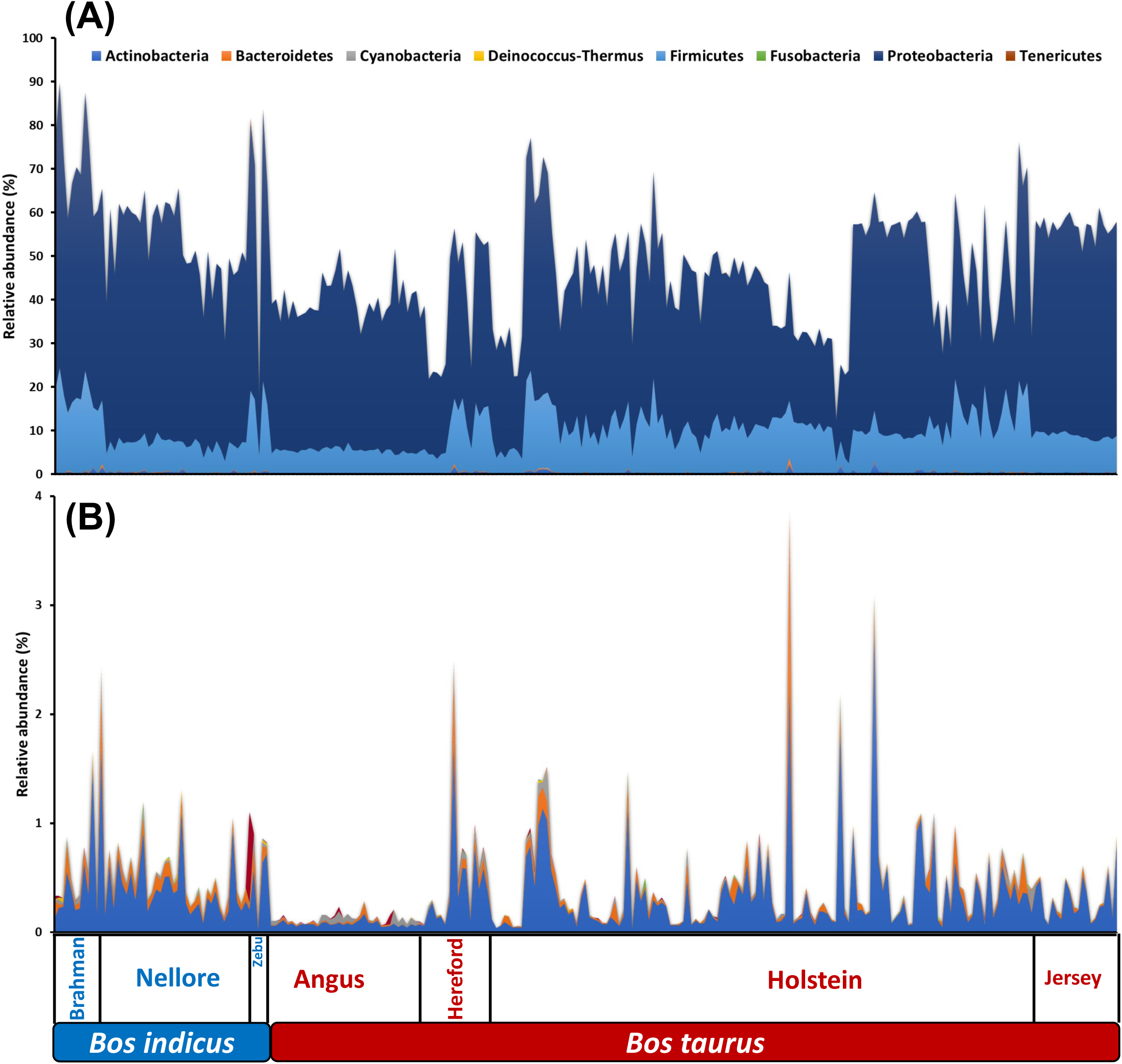
Relative abundance of core liver bacterial phyla (detected at least 50% of the meta-analyzed samples) across 251 liver samples. (A) All and (B) minor phyla (less than 1% relative abundance across all liver samples) were plotted separately.

Among the 39 core bacterial genera identified in the liver metatranscriptome, 21 genera belonged to the Proteobacteria phylum, while seven and nine genera were found within Actinobacteria and Firmicutes lineage, respectively (**Figure 2**). Two genera within Bacteroidetes were determined as core member in the liver microbiota. The relative abundance of general gut bacterial genera varies with that of their own habitat. The relative abundance of *Clostridium* was still dominant in the liver (6.30±0.04%) but that of omnipresent rumen bacterial genus *Prevotella* was quite limited (0.0037±0.0002%) with moderate occurrence rate in the liver (60.16%).

**Figure 2.**
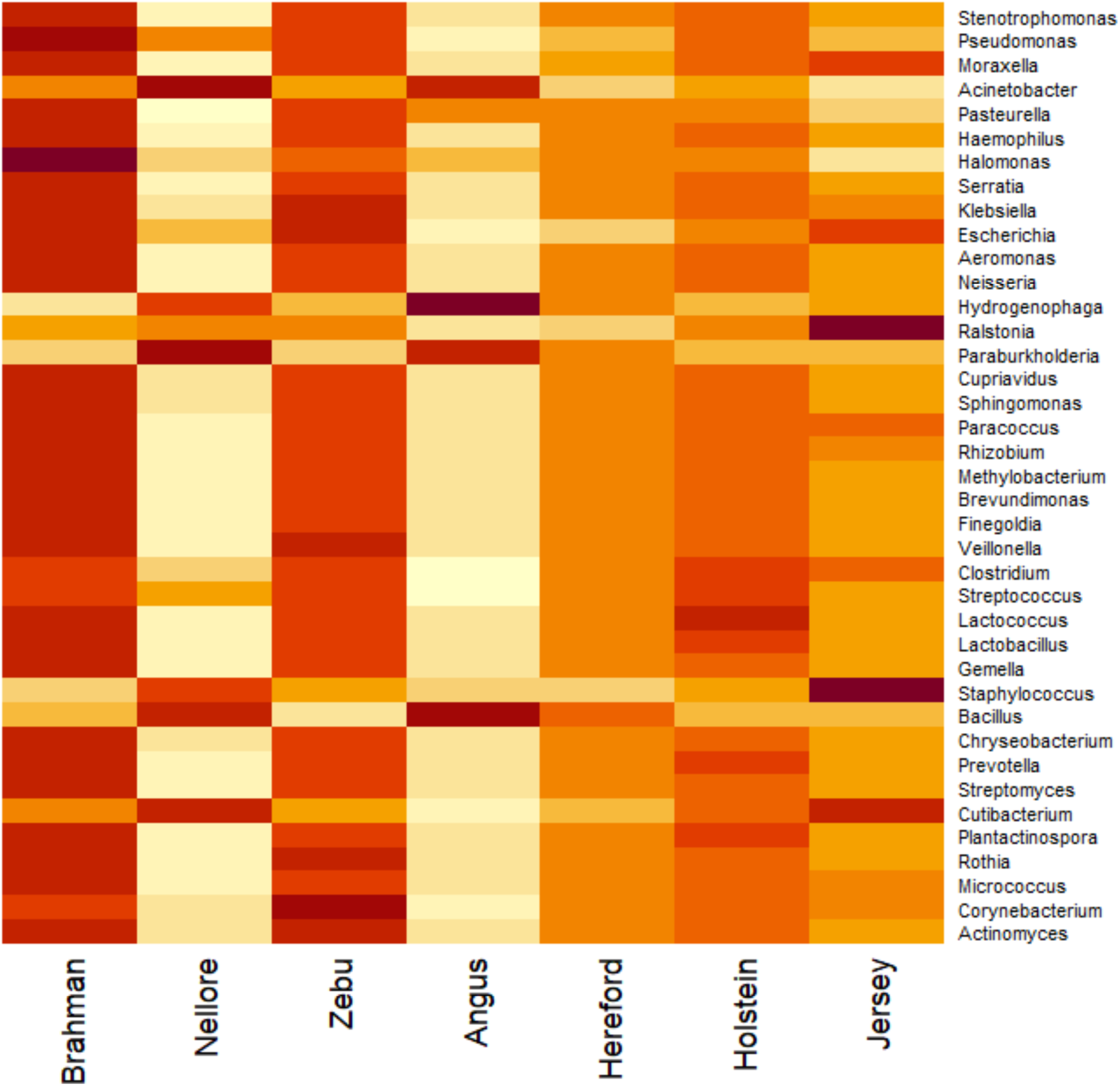
Relative abundance of core liver bacterial genera (detected at least 50% of the meta-analyzed samples) across 251 liver samples.

### Microbial and functional biomarkers associated with the ruminant species, breeds, and ages

By adjusting covariates commonly identified in the all 251 liver metatranscriptome samples, 6, 2, and 2 bacterial phyla were significantly associated with ruminant species, breed, and age (*Q* < 0.05), respectively (**Figure 3**). Tenericutes and its lower class, Mollicutes was associated with both the ruminant species and breed, negatively with *Bos taurus* and Nellore, respectively (*Q* = 0.0145 and *Q* = 0.0021). Veillonellaceae and its order and class level taxa, Negativicutes and Veillonellales, were negatively associated with *Bos taurus* (*Q* < 0.05). Flavobacteriaceae was the only taxon within Bacteroidetes showing significance in association with ruminant species. Bifidobacteriaceae and its order level taxon, Bifidobacteriales were positively associated with cattle age (*Q* = 0.0352). No core bacterial genera was significantly associated with any of those covariates used in this study. However, 5 and 7 minor bacterial taxa were significantly associated with ruminant species and breed, respectively (Table S2).

**Figure 3.**
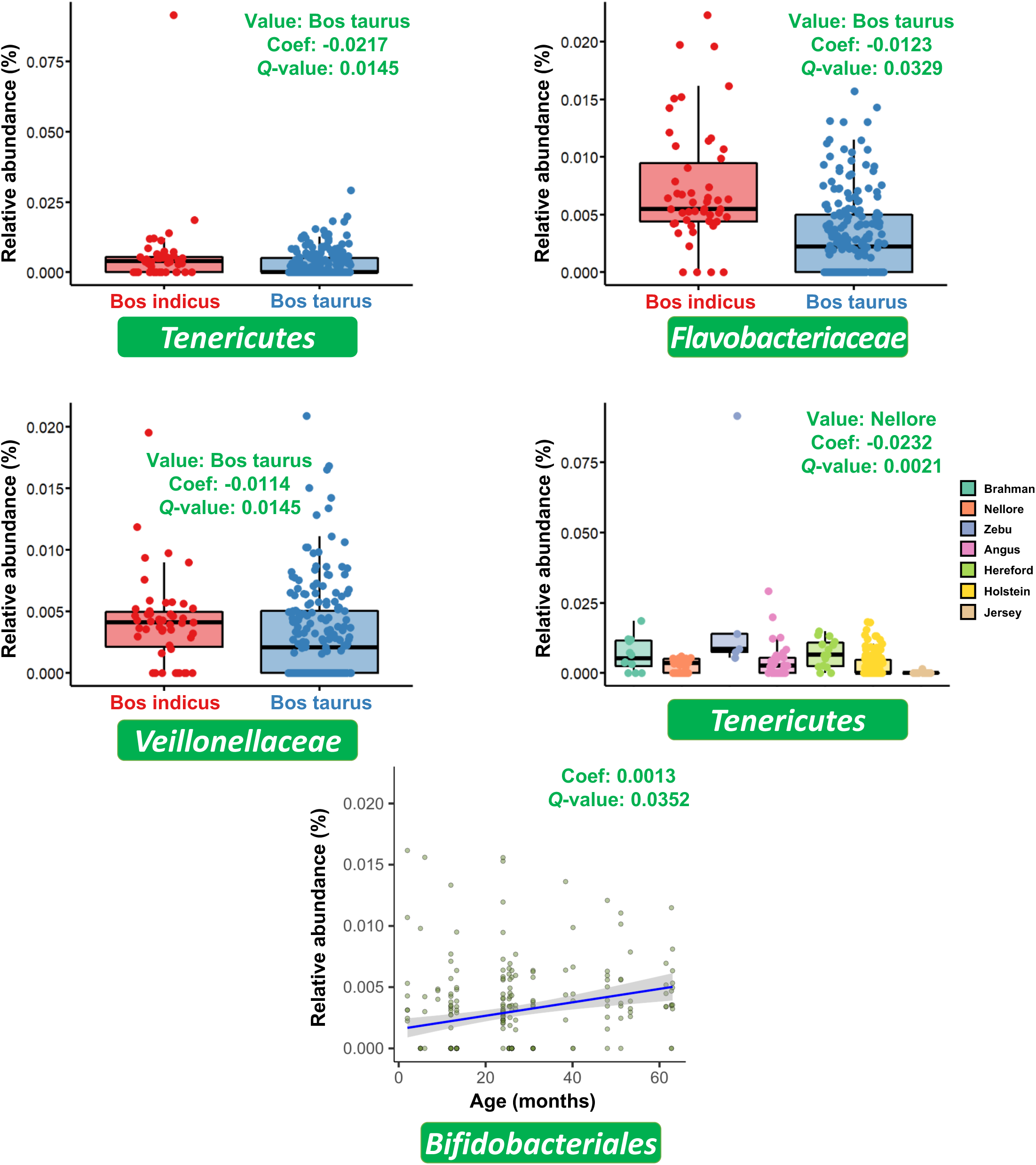
Cattle covariates (species, breeds, and age) associated liver bacterial taxa (*Q* < 0.05). Value, categorical variables associated with bacterial taxa; Coef, coefficient; *Q*-value, *P*-value corrected by the Benjamini-Hochberg method.

**Table 2.** Batch effect adjustment result using MMUPHin.

| Liver microbiota |  |  |  |  |  |  |  |
| --- | --- | --- | --- | --- | --- | --- | --- |
| Before adjustment |  | Df | SumsOfSqs | MeanSqs | F.Model | R2 | Pr(>F) |
|  | Label | 13 | 12.1332 | 0.9333 | 141.100 | 0.8813 | 1.00E-04 |
|  | Residuals | 247 | 1.6338 | 0.0066 | 0.1187 |  |  |
|  | Total | 260 | 13.767 | 1 |  |  |  |
| After adjustment |  | Df | SumsOfSqs | MeanSqs | F.Model | R2 | Pr(>F) |
|  | Label | 13 | 7.6948 | 0.5919 | 18.895 | 0.4986 | 1.00E-04 |
|  | Residuals | 247 | 7.7377 | 0.0313 | 0.5014 |  |  |
|  | Total | 260 | 15.4325 | 1 |  |  |  |
| Liver microbial functional features |  |  |  |  |  |  |  |
| Before adjustment |  | Df | SumsOfSqs | MeanSqs | F.Model | R2 | Pr(>F) |
|  | Label | 13 | 20.998 | 1.6153 | 24.189 | 0.5601 | 1.00E-04 |
|  | Residuals | 247 | 16.494 | 0.0668 | 0.43992 |  |  |
|  | Total | 260 | 37.492 | 1 |  |  |  |
| After adjustment |  | Df | SumsOfSqs | MeanSqs | F.Model | R2 | Pr(>F) |
|  | Label | 13 | 27.679 | 2.1292 | 14.358 | 0.4304 | 1.00E-04 |
|  | Residuals | 247 | 36.627 | 0.1483 | 0.5696 |  |  |
|  | Total | 260 | 64.306 | 1 |  |  |  |

Among the core-microbial functional features, represented by KEGG pathways, ko03060 (protein export) related to the active transport of protein by gram-negative bacteria was negatively associated with Hereford breed (*Q* = 0.0241) (**Figure 4**). Eight core KEGG pathways were either positively [ko03008 (Ribosome biogenesis in eukaryotes), ko03320 (PPAR signaling pathway), ko04626 (Plant-pathogen interaction), and ko04712 (Circadian rhythm - plant)] or negatively [ko03060, ko04962 (Vasopressin-regulated water reabsorption), ko05110 (*Vibrio cholerae* infection), and ko05146 (Amoebiasis)] associated with cattle age. Among the minor ones, 4 and 3 KEGG pathways were significantly associated either with ruminant breed and age, respectively (**Table 1**). No core and minor KEGG pathways were significantly associated with ruminant species.

**Figure 4.**
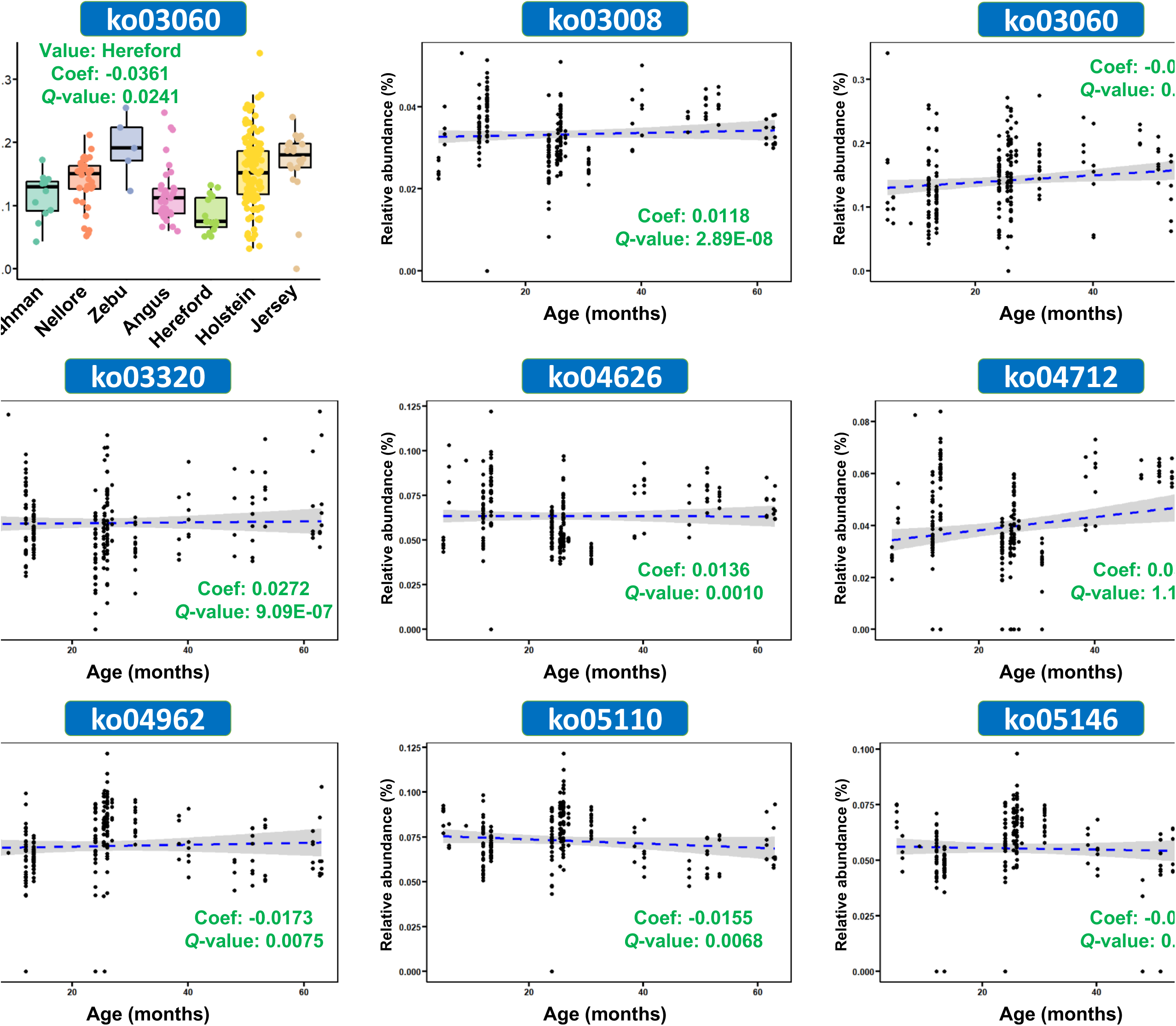
Cattle covariates (species, breeds, and age) associated liver microbial functions (KEGG pathways) (Q<0.05). Value, categorical variables associated with bacterial taxa; Coef, coefficient; *Q*-value, *P*-value corrected by the Benjamini-Hochberg method.

### Diversity measurements and overall bacterial and functional distribution in liver microbiome

Distribution of alpha-diversity measurements for both the bacterial genera abundance profiles and microbial functional features represented by KEGG orthologs show different distribution patterns across the chosen indices (Supplementary **Figure S1**). However, there were no alpha-diversity measurements from both bacterial and functional features significantly associated with any of the covariates.

Covariate-controlled, batch-effect adjustment resulted in reduced variability explained by study differences of about 38% and 13% for both the bacterial and functional abundance, respectively, but did not eliminate it in both the bacterial and functional abundance profiles (**Table 2**). PERMANOVA analysis showed that ruminant breed significantly affected both the core bacterial genera and liver functional features (**Figure 5 & Table 3**). Two different ruminant species had significantly different functional communities (adjusted *P*-value = 0.0032), but this was not derived from the core liver microbiota difference (adjusted *P*-value = 0.0565) (**Figure 5A**). Among the pairwise comparisons between different cattle breeds, Holstein and Hereford did not have different core liver bacterial communities (adjusted *P*-value = 0.3234) while Brahman and Jersey had similar functional communities with that of Holstein (adjusted *P*-value > 0.1) (Figure 5B). Angus and Brahman also had similar functional communities (adjusted *P*-value = 0.0798). Other comparisons for both the core bacterial and functional communities were all significantly different. Distribution of overall bacterial and functional feature communities by cattle age were also shown in **Figure 5C**. We compared the rRNA transcript abundance in the liver across the three groups of cattle, identifying 17 phyla with significantly different abundances (normalized-Read-count>2 and *P*-value <0.001) amongst them (**Figure 6**).

**Figure 5.**
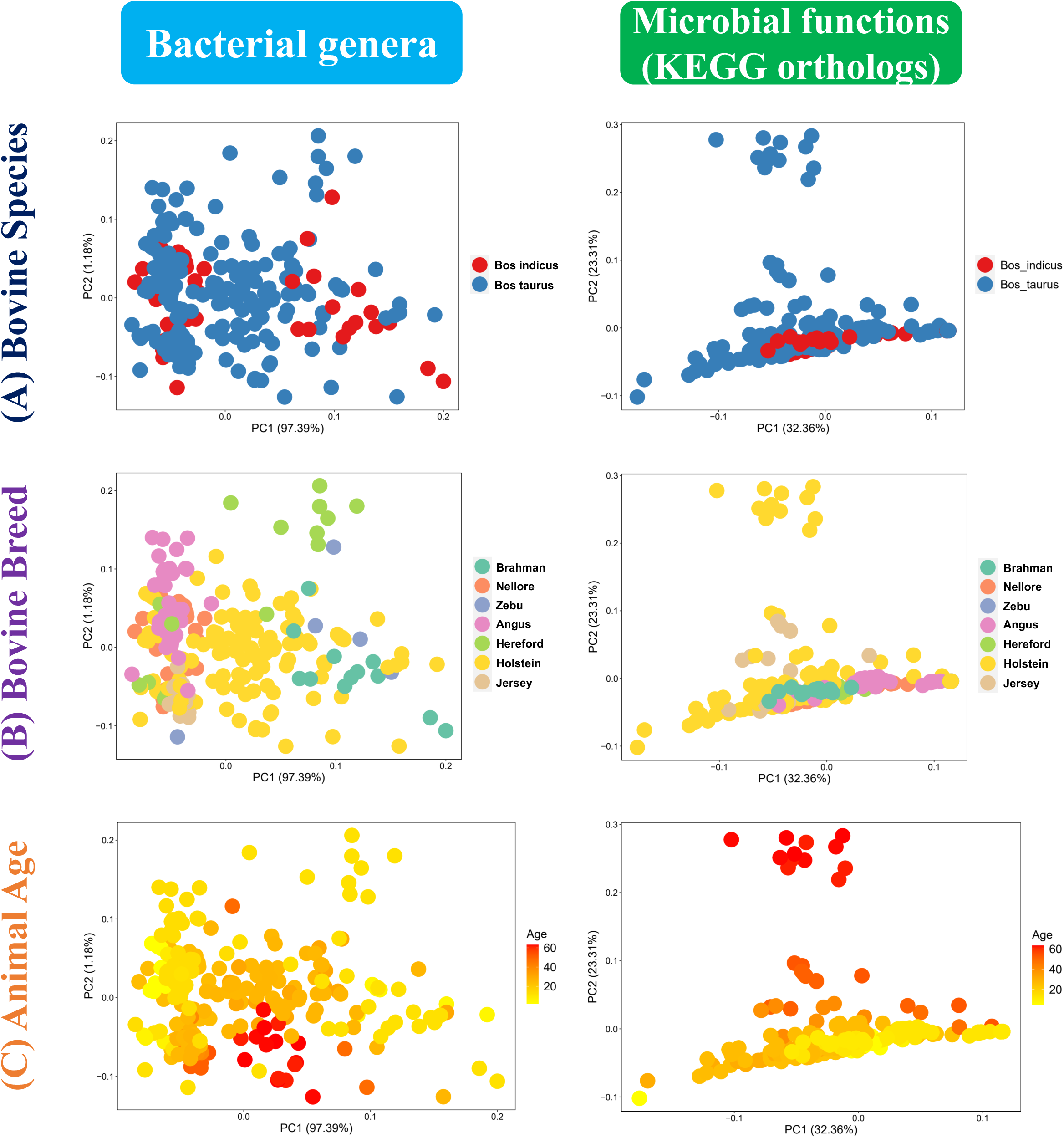
Principle components analysis plot showing the overall distribution of liver active microbiome represented by bacterial genera and KEGG ortholog profiles differed by (A) bovine species, (B) bovine breeds, and (C) animal ages.

**Figure 6.**
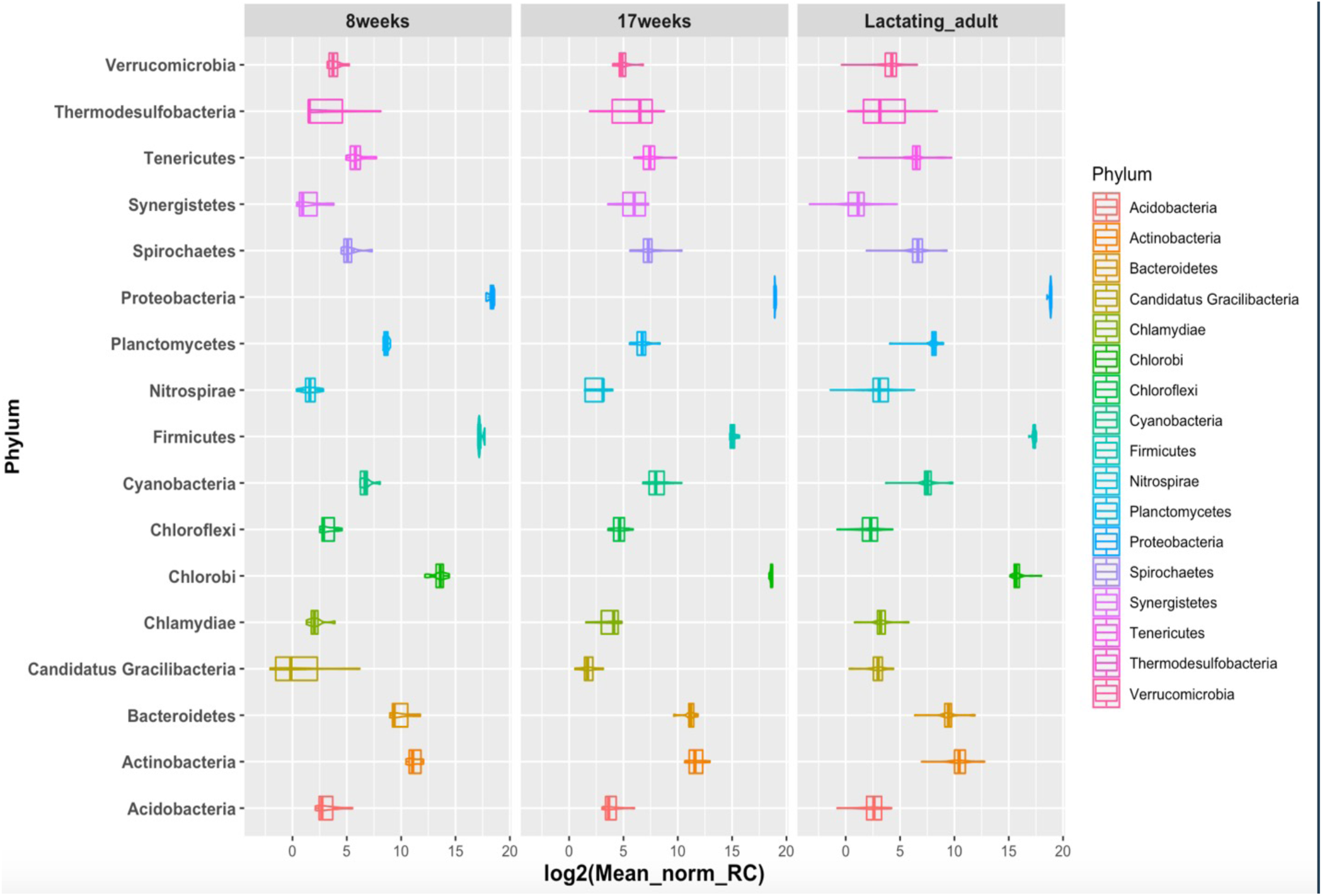
The genera with significantly different abundances in liver tissues across three age groups of cattle: 8-week, 17-week and lactating adult.

**Table 3.** Pairwise comparisons of Bray-Curtis distance matrices by different covariates in liver microbiome.

| Core liver microbiota (Genus level) |  |  |  |  |  |  |  |
| --- | --- | --- | --- | --- | --- | --- | --- |
|  | Brahman | Nellore | Zebu | Angus | Hereford | Holstein | Jersey |
| Brahman |  | 0.0021 | 1 | 0.0021 | 0.0168 | 0.0021 | 0.0021 |
| Nellore | 0.0021 |  | 0.0042 | 0.0021 | 0.0021 | 0.0021 | 0.0021 |
| Zebu | 1 | 0.0042 |  | 0.0021 | 1 | 1 | 0.0063 |
| Angus | 0.0021 | 0.0021 | 0.0021 |  | 0.0021 | 0.0021 | 0.0021 |
| Hereford | 0.0168 | 0.0021 | 1 | 0.0021 |  | 0.3234 | 0.0021 |
| Holstein | 0.0021 | 0.0021 | 1 | 0.0021 | 0.3234 |  | 0.0021 |
| Jersey | 0.0021 | 0.0021 | 0.0063 | 0.0021 | 0.0021 | 0.0021 |  |
| Core liver functional features (KEGG-orthologs) |  |  |  |  |  |  |  |
|  | Brahman | Nellore | Zebu | Angus | Hereford | Holstein | Jersey |
| Brahman |  | 0.0042 | 0.0021 | 0.0798 | 0.0042 | 0.4893 | 0.0021 |
| Nellore | 0.0042 |  | 0.0021 | 0.0021 | 0.0021 | 0.0021 | 0.0021 |
| Zebu | 0.0021 | 0.0021 |  | 0.0042 | 1 | 0.0021 | 0.0021 |
| Angus | 0.0798 | 0.0021 | 0.0042 |  | 0.0021 | 0.0021 | 0.0042 |
| Hereford | 0.0042 | 0.0021 | 1 | 0.0021 |  | 0.0021 | 0.0021 |
| Holstein | 0.4893 | 0.0021 | 0.0021 | 0.0021 | 0.0021 |  | 0.1344 |
| Jersey | 0.0021 | 0.0021 | 0.0021 | 0.0042 | 0.0021 | 0.1344 |  |

| Core liver microbiota (Genus level – upper right) |  |  |  |  |  |  |  |
| --- | --- | --- | --- | --- | --- | --- | --- |
|  | Brahman | Nellore | Zebu | Angus | Hereford | Holstein | Jersey |
| Brahman |  | 0.0021 | 1 | 0.0021 | 0.0168 | 0.0021 | 0.0021 |
| Nellore | 0.0042 |  | 0.0042 | 0.0021 | 0.0021 | 0.0021 | 0.0021 |
| Zebu | 0.0021 | 0.0021 |  | 0.0021 | 1 | 1 | 0.0063 |
| Angus | 0.0798 | 0.0021 | 0.0042 |  | 0.0021 | 0.0021 | 0.0021 |
| Hereford | 0.0042 | 0.0021 | 1 | 0.0021 |  | 0.3234 | 0.0021 |
| Holstein | 0.4893 | 0.0021 | 0.0021 | 0.0021 | 0.0021 |  | 0.0021 |
| Jersey | 0.0021 | 0.0021 | 0.0021 | 0.0042 | 0.0021 | 0.1344 |  |
| Core liver functional features (KEGG-orthologs – lower left) |  |  |  |  |  |  |  |

### Cattle liver resistome

Heatmap and hierarchical clustering of mean RPKM profiles of detected ARGs from cattle liver metatranscriptome showed clusters were not associated with ruminant species but with breed (**Figure 7**). Beta-lactam resistance was the most represented ARG in cattle liver regardless of ruminant breed and reference databases. Cattle breeds associated with antimicrobial resistance genes including biguanide and elfamycins resistance (**Table 4**). Biguanide resistance was negatively associated with three *Bos taurus* breeds, Holstein, Hereford, and Jersey. Elfamycin was positively associated with Hereford breed. No ruminant species and age effects were shown on the ARGs distribution in liver.

**Figure 7.**
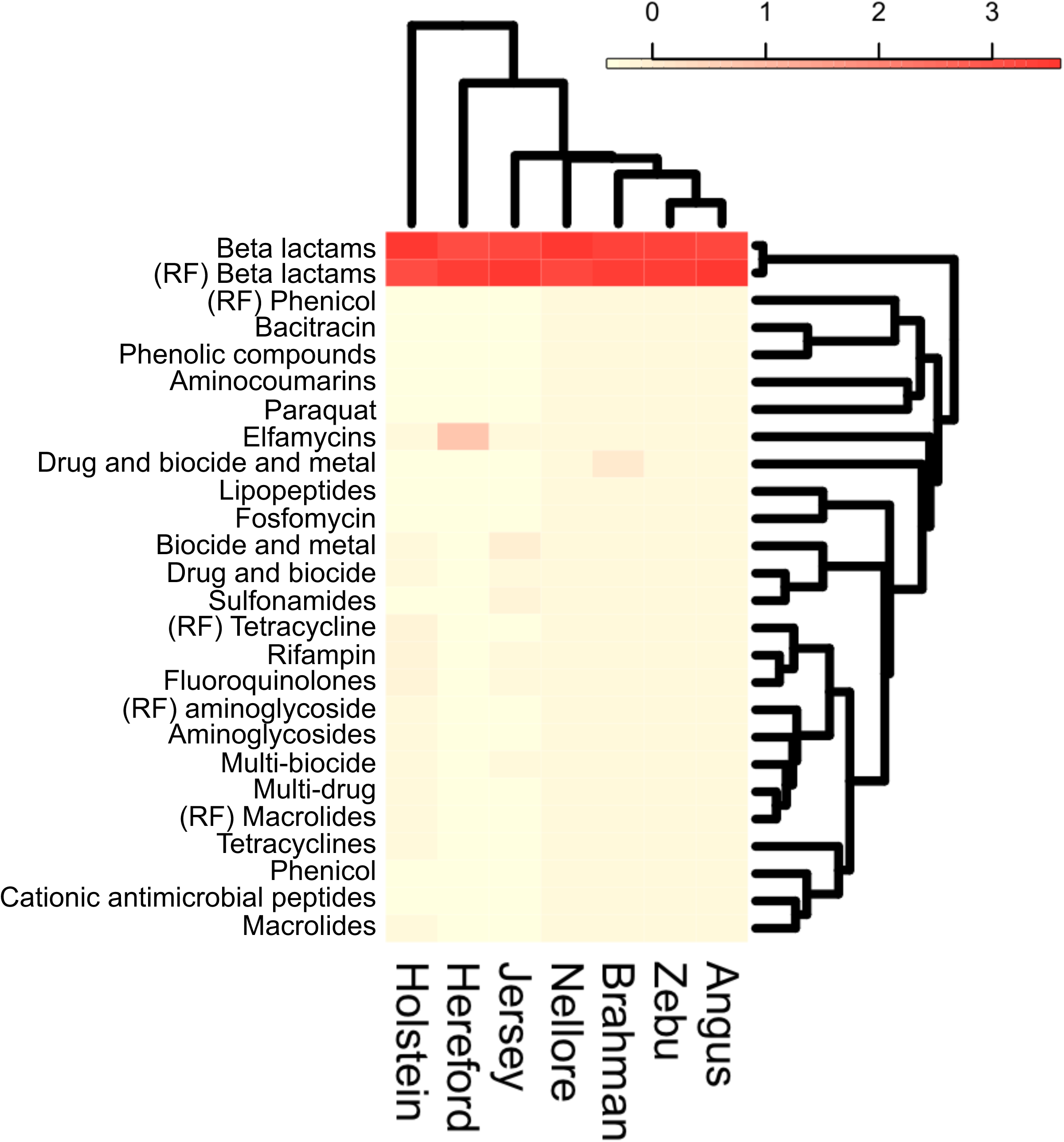
Heatmap and hierarchical dendrogram of antimicrobial-resistance gene RPKM profiles across ruminant breeds. Red represents the highest average RPKM and yellow represents the lowest value for the ruminant breeds. RF, ResFinder annotated ARGs. Others were annotated based on the MEGARes databases.

**Table 4.** Significant associations with meta-analyzed covariates among ARGs using MaAsLin2.

| Feature | Value | Coefficient | Standard error | No. of samples | No. of samples not zero | *P*-value | *Q*-value |
| --- | --- | --- | --- | --- | --- | --- | --- |
| Cattle breeds associated |  |  |  |  |  |  |  |
| Biguanide resistance | Hereford | -0.0009 | 0.0002 | 251 | 8 | 2.81E-05 | 0.0026 |
|  | Holstein | -0.0009 | 0.0002 | 251 | 8 | 1.25E-05 | 0.0021 |
|  | Jersey | -0.0009 | 0.0002 | 251 | 8 | 0.0007 | 0.0456 |
| Elfamycins | Hereford | 0.3047 | 0.0654 | 251 | 123 | 5.27E-06 | 0.0021 |

## Discussion

### Core microbial taxa identified in the liver

While there has been considerable work focusing on the liver of diseased cattle, our study is primarily a survey of cattle liver samples without obvious liver disease. Among the core phyla identified in our study, four of them (Actinobacteria, Bacteroidetes, Firmicutes, and Fusobacteria) were identified as dominant phyla in the abscessed liver in feedlot cattle using a targeted, 16S rRNA amplicon sequencing approach (Weinroth, Carlson et al. 2017). Three studies using DNA-based 16S rRNA amplicon sequencing have identified Fusobacteria, Bacteroidetes and Proteobacteria as the most abundant phyla in abscessed cattle liver (Amachawadi, Tom et al. 2021). The Fusobacteria has been reported as the causal microbe of liver abscess in several studies and is the most common cause of liver abscesses (Calkins and Dewey 1968, Narayanan, Nagaraja et al. 1997). *Fusobacterium* is an anaerobic gram-negative bacillus that includes 13 species. *Fusobacterium necrophorum* is a normal habitant of the alimentary tract of animals (Tan, Nagaraja et al. 1996). Two sub-species of *Fusobacterium necrophorum* have been recognized: *F. necrophorum,* subsp. *Necrophorum* (biotype A) and subsp. *Funduliforme* (biotype B). The biotype A is more virulent since it’s more frequently isolated from infections than the biotype B. The biotype A is considered an opportunistic pathogen in a variety of animals. Bovine liver abscesses and foot rot are among the infections caused primarily by the *Fusobacterium necrophorum* (Nagaraja and Lechtenberg 2007). The translocation of the *Fusobacteria* from the rumen into the portal circulatory system is considered as the mechanisms of bovine liver abscess, which is often sequela to ruminal acidosis in cattle fed a highly fermentable diet (Amachawadi, Purvis et al. 2017).

Culture based methods using abscessed liver samples from both beef and dairy cattle indicated that *Trueperella pyogenes* and *F. necrophorum* are present in all of the studied samples (Amachawadi, Purvis et al. 2017) and the concentration of *T. pyogenes* is higher in crossbred cattle compared to Holstein steers. *T. pyogenes* is part of the skin biota and mucous membrane of the upper respiratory and gastrointestinal tracts of animals (Queen, Ward et al. 1994, Silva, Gaivao et al. 2008). Specifically, this bacterium was isolated from the bovine rumen epithelial (Narayanan, Nagaraja et al. 1998) and from the udder of clinically healthy cows (Spittel and Hoedemaker 2012). Though *T. pyogenes* is a commensal bacterium, it is a common opportunistic pathogen, which causes purulent or necrotic lesions in several different tissues, including uterus (Amos, Healey et al. 2014, Bicalho, Lima et al. 2016), mammary gland (Madsen, Aalbaek et al. 1992, Rezanejad, Karimi et al. 2019), liver (Tadepalli, Narayanan et al. 2009), and hoof (Kontturi, Junni et al. 2019). Stotz and coauthors (Stotz, Henry et al. 2021) reported Tenericutes was identified as a dominant phylum second to Fusobacteria and Bacteroidetes in severely abscessed liver samples. In the same study, Cyanobacteria was identified as the dominant phyla in non-abscessed and scarred liver samples. Some studies indicated that some pathogenic bacteria (e.g., *Mycobacterium tuberculosis*, *Salmonella typhimurium*, and *Staphylococcus aureus*) have developed mechanisms to evade host defense in the liver and may later cause tissue damage and death (Engele, Stossel et al. 2002).

The predominance of *Proteobacteria* is identified in our analysis. Several pathogenic genera within this *Proteobacteria* have been reported. The commonly held perception is that a pathological state or infection of the liver is necessarily associated with the liver microbes. They include *Escherichia*, *Salmonella*, *Vibrio*, *Yersinia*, *Legionella*, and many others. Their outer membrane mainly composed of lipopolysaccharides (LPS), which is widely regarded as a potent activator of monocytes/macrophages (Takashiba, Van Dyke et al. 1999). By triggering the release of a vast number of inflammatory cytokines in various cell types, LPS can cause significant inflammation (Ngkelo, Meja et al. 2012). Quite interestingly, Proteobacteria was identified as one of the predominant phylum in non-abscessed livers (Stotz, Henry et al. 2021), consistent with the report that this phylum is dominant in the rumen (Mann, Wetzels et al. 2018).

We discovered abundant microbial taxa in the liver that varied by age, species and developmental stage. The differential abundance of the bacterial phyla might be leveraged as potential biomarkers for the pathogenesis of the liver.

### Cattle species, breed, and age effects on core-liver taxa and microbial functions

We observed significant impact of the species and breed on the variation of core-liver taxa and microbial functions. It’s known that the incidence of liver abscesses is highly variable and the cattle type is one of several factors influencing the development of liver abscesses (Nagaraja and Chengappa 1998, Amachawadi, Purvis et al. 2017). Holstein steers tend to have a higher prevalence and severity of liver abscesses than the beef cattle breeds (Amachawadi and Nagaraja 2016), potentially due to the increased days on high-grain finishing diet and higher intake (Duff and McMurphy 2007). Another factor that potentially contributes to the species-and breed-specific core-liver microbial taxa and functions is the dietary characteristics. Specifically, the amount and type of forage include in the finishing diet have been associated with the incidence of liver abscess in cattle. Utley and coauthors reported that the physical characteristics of the forage tend to impact the incidence of liver abscess (Utley, Hellwig et al. 1973), in which a higher rate of liver abscess was observed in the steers fed ground or pelleted peanut hulls.

We identified several KEGG pathways with significant association with cattle age. They include the positively correlated: ko03008 (Ribosome biogenesis in eukaryotes), ko03320 (PPAR signaling pathway), ko04626 (Plant-pathogen interaction), and ko04712 (Circadian rhythm - plant), and negatively correlated: ko03060, ko04962 (Vasopressin-regulated water reabsorption), ko05110 (Vibrio cholerae infection), and ko05146 (Amoebiasis). This suggest that the liver microbial community might evolve as the host ages. Thus, there is a need to investigate the liver microbiome at different developmental stages of cattle’s life to identify how the microbes change abundances with animal age.

### Microbial circulation in cattle blood before its colonization in the liver

Liver abscess has an incidence rate of 12-32% in grain-finished feedlot cattle (Nagaraja, Laudert et al. 1996, Radostits, Gay et al. 2000) and it’s typically observed at the time of culling. Despite significant economic loss due to liver abscess, there is currently no effective method available for early detection prior to culling. The liver sits at the junction between the GIT and systemic circulation, in part due to its intimate proximity to the GIT via circulatory loop where the liver receives ∼80% of its blood supply from the GIT via the portal vein (Lomax and Baird, 1983). Gut microbiota and bacterial products can directly and indirectly affect the liver, leading to the development of a wide range of hepatic disorders (Cani and Delzenne 2009, Duseja and Chawla 2014) through multiple mechanisms, including increased GIT permeability (i.e., leaky gut), chronic systemic inflammation, and production of short chain fatty acids (SCFAs) that elicit changes in liver metabolism (Schwenger, Clermont-Dejean et al. 2019). Before the microbiome colonizes the liver, it’s reasonable to hypothesize that the commensal or pathogenic microbes are in the blood and could potentially enter systemic circulation. Studies evaluating the hypothesis of the existence of a systematic blood microbiome are needed to evaluate potential functionality in healthy and diseased states and to develop early, microbe-based diagnostic biomarkers for the early detection of liver or systemic diseases.

## Conclusions

While microbiota have long been known to contribute to pathologies such as liver abscesses in feedlot cattle, this is the first study in which a core microbiome has been described in cattle without overt liver disease. These results provide foundational knowledge of the core liver microbiome and microbial functions as identified by a whole transcriptome sequencing approach in cattle of different species, breed, and age. Several phyla identified in our study have corroborative evidence from previously published studies using DNA-based, 16S rRNA amplicon sequencing. However, several core phyla identified in our study were not previously reported, highlighting the improved sensitivity or ability in detecting microbes by RNA-over DNA-based methods. Furthering the application of transcriptome sequencing-based analysis may additionally help identify the active microbes in the liver and further facilitate the blood or liver microbe biomarkers discovery for early diagnosis and prevention of liver diseases in cattle.

## List of abbreviation

GIT: gastrointestinal tract
SRA: Sequence Read Archive
NCBI: National Center for Biotechnology Information
KO: KEGG orthologs
KAAS: KEGG Automatic Annotation Server
ARG: antimicrobial-resistance gene
RPKM: Reads per kilo base per million mapped reads
PCA: principal component analysis

## Ethics declarations

### Ethics approval and consent to participate

Not applicable.

### Consent for publication

Not applicable.

### Availability of data and material

The accession numbers for all the publicly available data are listed in this manuscript. For unpublished data, it’s available upon reasonable request.

### Competing interests

The authors declare no conflicts of interest associated with this study.

### Funding

This work is funded by USDA Agricultural Research Services, funding number: 5090-31000-026-000D 02. TP is supported by a fellowship from Oak Ridge Institute for Science and Education. This research used resources provided by the SCINet project of the USDA Agricultural Research Service, ARS project number 0500-00093-001-00-D.

#### Authors’ contributions

WL conceived the study, performed the data collection; WL, GZ, and TP designed the data analysis plan. TP and WL performed the bioinformatic analysis. TP and WL wrote the manuscript. TP, WL and GZ edited the manuscript. All the authors read and approved the final version of the manuscript.

## Acknowledgements

We thank the colleagues at US Dairy Forage Research Center for animal and farm support. We thank the members at WL’s laboratory for collecting raw tissues and perform RNA sequencing experiments.

## Competing interests

The authors declare no conflicts of interest associated with this study.

## Figure legends

**Supplementary Figure S1.**
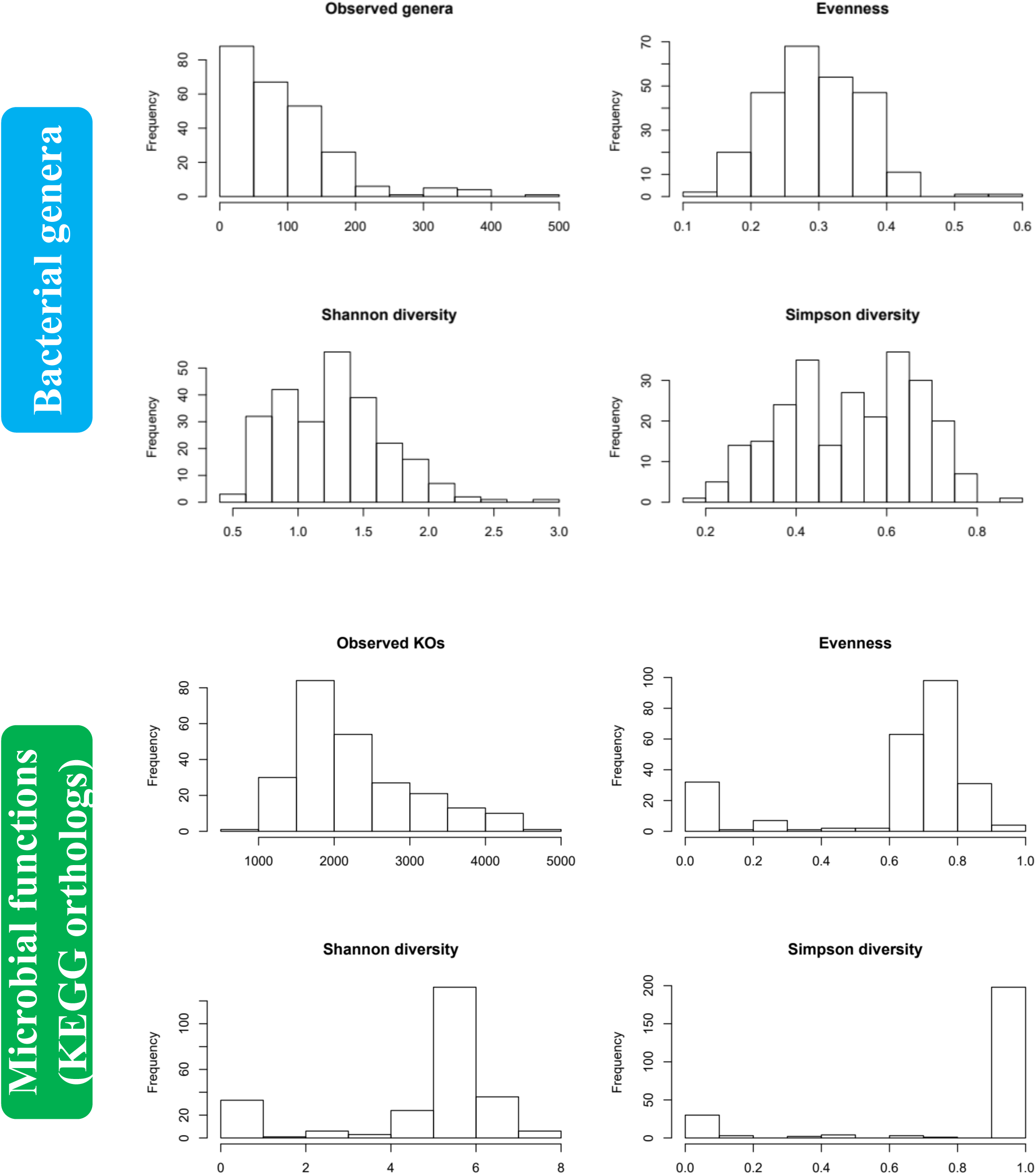
Alpha-diversity histogram.

**Supplementary Table S1.** Dataset for the liver whole transcriptome.

| **NCBI Accession #** | **Cattle breed** | **Age** | **Sex** | **Total # of samples** |
| --- | --- | --- | --- | --- |
| PRJEB8831 | Nellore cattle | Adult | Male | 16 |
| PRJEB27337 | Nellore cattle | Adult | Male | 18 |
| PRJNA263327 | Holstein | Adult (late pregnancy to mid lactation) | Female | 48 |
| PRJNA303131 | Holstein | Adult, 3 wk antepartum and 3 wk postpartum | Female | 17 |
| PRJNA312148 | Hereford, Polish Holstein | 6m to 12 m of age | Male | 11 |
| PRJNA357463 | Holstein and Jersey | Adults with 1-3 parity | Female | 38 |
| PRJNA392196 | Holstein | Adult (lactation) | Female | 10 |
| PRJNA393239 | Angus | Adult | Male | 37 |
| PRJNA436715 | Brahman | Pre- and post-puberty | Female | 12 |
| PRJNA508221 | Hereford | Adult | Male | 10 |
| PRJNA526337 | Zebu (Bos indicus) and Holstein | Adult | Female | 10 |
| PRJNA597813 | Holstein | Young calf | Male | 6 |
| PRJNA624810 | Holstein | Adult (lactation) | Female | 10 |
| in-house study | Holstein | 8 weeks of age | Male | 8 |

**Supplementary Table S2.** Significant associations with meta-analyzed covariates among minor bacterial taxa using MaAsLin2.

| Feature | Value | Coefficient | Standard error | No. of samples | No. of samples not zero | P-value | Q-value | Taxonomy lineage |
| --- | --- | --- | --- | --- | --- | --- | --- | --- |
| Cattle species associated |  |  |  |  |  |  |  |  |
| Beijerinckiaceae | Bos taurus | 0.0013 | 0.0002 | 251 | 26 | 4.96E-08 | 5.62E-05 | p__Proteobacteria\|c__Alphaproteobacteria\|o__Rhizobiales\|f__Beijerinckiaceae |
| Rhodobacterales | Bos taurus | -0.0086 | 0.0018 | 251 | 207 | 3.21E-06 | 0.0020 | p__Proteobacteria\|c__Alphaproteobacteria\|o__Rhodobacterales |
| Rhodobacteraceae | Bos taurus | -0.0085 | 0.0018 | 251 | 207 | 3.02E-06 | 0.0020 | p__Proteobacteria\|c__Alphaproteobacteria\|o__Rhodobacterales\|f__Rhodobacteraceae |
| Melaminivora | Bos taurus | -0.0011 | 0.0002 | 251 | 39 | 1.32E-06 | 0.0010 | p__Proteobacteria\|c__Betaproteobacteria\|o__Burkholderiales\|f__Comamonadaceae\|g__Melaminivora |
| Candidatus Carsonella | Bos taurus | 0.0032 | 0.0005 | 251 | 49 | 2.06E-10 | 8.16E-07 | p__Proteobacteria\|c__Gammaproteobacteria\|o__Oceanospirillales\|f__Halomonadaceae\|g__Candidatus Carsonella |
| Cattle breeds associated |  |  |  |  |  |  |  |  |
| Acidobacteriales | Brahman | 0.0037 | 0.0010 | 251 | 32 | 0.0002 | 0.0493 | p__Acidobacteria\|c__Acidobacteriia\|o__Acidobacteriales |
| Acidobacteriaceae | Brahman | 0.0037 | 0.0010 | 251 | 32 | 0.0002 | 0.0493 | p__Acidobacteria\|c__Acidobacteriia\|o__Acidobacteriales\|f__Acidobacteriaceae |
| Hathewayia | Brahman | 0.0057 | 0.0008 | 251 | 42 | 6.88E-06 | 0.0036 | p__Firmicutes\|c__Clostridia\|o__Clostridiales\|f__Clostridiaceae\|g__Hathewayia |
| Beijerinckiaceae | Hereford | -0.0013 | 0.0003 | 251 | 26 | 2.16E-05 | 0.0107 | p__Proteobacteria\|c__Alphaproteobacteria\|o__Rhizobiales\|f__Beijerinckiaceae |
|  | Holstein | -0.0010 | 0.0003 | 251 | 26 | 0.0001 | 0.0410 |  |
| Chelatococcaceae | Brahman | 0.0022 | 0.0005 | 251 | 27 | 6.00E-05 | 0.0236 | p__Proteobacteria\|c__Alphaproteobacteria\|o__Rhizobiales\|f__Chelatococcaceae |
| Chelatococcus | Brahman | 0.0022 | 0.0005 | 251 | 27 | 6.00E-05 | 0.0236 | p__Proteobacteria\|c__Alphaproteobacteria\|o__Rhizobiales\|f__Chelatococcaceae\|g__Chelatococcus |
| Candidatus Carsonella | Holstein | -0.0034 | 0.0005 | 251 | 49 | 2.86E-09 | 4.57E-06 | p__Proteobacteria\|c__Gammaproteobacteria\|o__Oceanospirillales\|f__Halomonadaceae\|g__Candidatus Carsonella |
|  | Hereford | -0.0024 | 0.0006 | 251 | 49 | 8.74E-05 | 0.0288 |  |
|  | Jersey | -0.0029 | 0.0007 | 251 | 49 | 9.45E-05 | 0.0288 |  |

## Notes

### Competing Interest Statement

The authors have declared no competing interest.

